# Characterizing neural phase-space trajectories via Principal Louvain Clustering

**DOI:** 10.1101/2021.03.08.434355

**Authors:** Mark M. Dekker, Arthur S. C. França, Debabrata Panja, Michael X Cohen

## Abstract

**Background:** With the growing size and richness of neuroscience datasets in terms of dimension, volume, and resolution, identifying spatiotemporal patterns in those datasets is increasingly important. Multivariate dimension-reduction methods are particularly adept at addressing these challenges.

**New Method:** In this paper, we propose a novel method, which we refer to as Principal Louvain Clustering (PLC), to identify clusters in a low-dimensional data subspace, based on time-varying trajectories of spectral dynamics across multisite local field potential (LFP) recordings in awake behaving mice. Data were recorded from prefrontal cortex, hippocampus, and parietal cortex in eleven mice while they explored novel and familiar environments.

**Results:** PLC-identified subspaces and clusters showed high consistency across animals, and were modulated by the animals’ ongoing behavior.

**Conclusions:** PLC adds to an important growing literature on methods for characterizing dynamics in high-dimensional datasets, using a smaller number of parameters. The method is also applicable to other kinds of datasets, such as EEG or MEG.

## 1. Introduction

The dimensionality of neuroscience measurements has increased drastically over the past few decades [1, 2], meaning that neuroscientists are now capable of recording simultaneous activity of dozens, hundreds, and even tens of thousands of neurons. Larger and richer datasets provide new opportunities for hypothesis-testing and exploratory discovery, but also challenges in conceptualizing and characterizing the multivariate signals.

Most traditional data analysis methods in neuroscience are univariate or mass-univariate, such as spike counts (in single-unit studies) or spectral power (in LFP studies). Univariate methods have been and remain the backbone of neuroscience data analysis; however, larger-scale datasets might benefit from multivariate analyses that can identify patterns embedded in population activity [3, 4].

Multivariate dimension-reduction methods seek to identify a low-dimensional subspace in which the most relevant activity patterns exist [5, 6, 7]. Such methods can identify meaningful structures that are embedded in the patterns of correlations across the data, which might be undetectable when considering the activity of individual neurons or LFP channels. This is due to neural activity being spatiotemporally synchronized across neurons and circuits, and across multiple spatial scales [8, 9].

However, neural data are not static inside those dimensions; they ebb and flow over time with cognitive/behavioral operations, and internal brain states [10, 11, 12]. Characterizing the time-varying trajectories inside these subspaces is often done visually, which is feasible only in 2 or 3 dimensions.

Here, we introduce a new method for clustering and characterizing trajectories of low-dimensional subspace neural dynamics. The method is based on a combination of principal components analysis (PCA) and Louvain clustering [13], and involves identifying discrete pockets in the principal component (PC) space in which the neural trajectory remains roughly stationary for some period of time. We therefore term this method PLC, for Principal Louvain Clustering. Louvain clustering [14] is a modularity optimization algorithm applicable to weighted graphs, and has been used to characterize spatiotemporal dynamics of railway traffic [13], clusters in air transport networks [15], diplomatic structures in formal alliance data [16] and atmospheric states [17].

We modified the Louvain clustering method to suit multisite LFP data recorded simultaneously from the prefrontal cortex, parietal cortex, and hippocampus, of awake behaving mice during novelty exploration. A spectral decomposition of the LFP via a 1*/ f* -removed short-time Fourier transform was applied, and a six-dimensional (6D) PC space was constructed based on the largest two components from each region. Each time index during the recording is a “brain state point” in this space, and thus successive time points create a trajectory over time. The Louvain clustering method identifies spatial clusters in which these trajectories remain roughly stationary for a period of time *τ*. Finally, characteristics of these clusters can be quantified and linked to the mice’ behavior during the novelty exploration task.

## 2. Materials and methods

### 2.1. Animals, electrodes, and behavioral task

Data are from 11 Black57 background male mice. Non-overlapping results from some of these data have been published elsewhere [18, 19]. The mice had free access to food and water. All experiments were approved by the Centrale Commissie Dierproeven (CCD), and the surgeries and experiments were conducted according to approved indications of the local Radboud University Medical Centre animal welfare body (Approval number 2016-0079).

Custom-designed and self-made electrode arrays were constructed to target three different regions of the mouse brain: the CA1 region of the hippocampus (HIP; 8 electrodes), parietal cortex (PAR; 8 electrodes), and the medial prefrontal cortex (PFC; 16 electrodes). Details of the arrays and the manufacturing process are available in [20].

For surgery, 10-16 week-old mice were anesthetized with isoflurane (induction at 5% isoflurane in 0.5L/min O_2_; maintenance at 1-2% in .5L/min O_2_). Mice were fixed in a Neurostar Stereotaxic frame. After shaving, the skin was disinfected with ethanol (70%). Local anesthetic xylocaine (2%, adrenaline 1:200,000 [AstraZeneca]) was injected subcutaneously at the incision site before exposing the skull. Peroxide (10-20% H_2_O_2_; [Sigma]) was applied to the skull with a cotton swab for cleaning and visualization of bregma and lambda. Electrodes and screws were fixed onto the skull with dental cement (Super-Bond C&B). Approximately 40 minutes prior to the end of the surgery, saline and analgesic (carprofen injected subcutaneous 2.5 mg/Kg) were injected to facilitate recovery.

After the experiments, the mice were euthanized for post-mortem histological confirmation of electrode location. The electrodes in PFC were distributed across anterior cingulate and secondary motor cortex. The PAR electrodes were placed among layers 2 to 5. HIP electrodes were located in the CA1 region, spread in different mice between *stratum pyramidale* and *stratum lacunosum-moleculare*.

Note that although the anatomical targets were the same in all mice, we do not assume that the electrodes have a one-to-one correspondence across mice (e.g., electrode #*x* in mouse 1 may not be not in the same functional location as electrode #*x* in mouse 2).

During the recordings, the mice performed a novelty-learning task as depicted in Figure 1a. For 10 minutes, they were placed in an arena with two objects that they could explore. Objects were every-day items such as a coffee mug or bath toy. This phase is called the “training phase”. After a 60-minute break, they were placed back in the arena for another 10 minutes, and one of the objects was replaced by a new object (orange circle in Figure 1a). In between these recording sessions, mice were placed in their home cage; data from those periods are not reported here.

**Figure 1.**
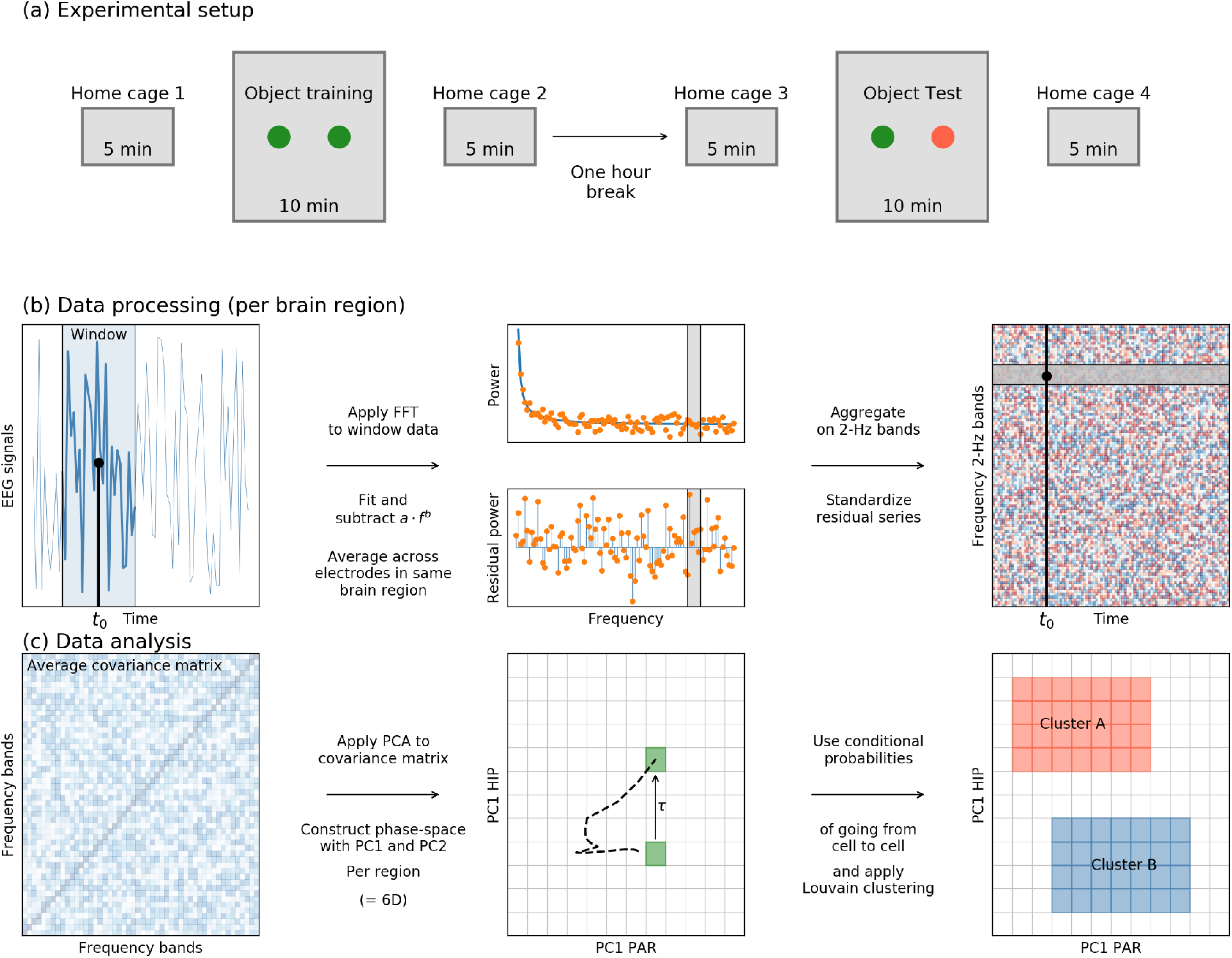
**Panel (a)**: the experimental setup. Each of 11 mice was held in a home cage for 5 minutes, and then was placed in an arena with two unknown objects (green dots) for 10 minutes. We refer to this as the *training phase*. After this, the mouse was returned to its home cage and was given a break for one hour. Then, the process was repeated except that one of the old objects was replaced with a new one (orange circle). We refer to this as the *test phase*. During both the training and test phases, the mouse’s LFP signals in the hippocampus, the prefrontal cortex and the parietal cortex were monitored by 32 electrodes. **Panel (b)**: The raw LFP signals were binned into a window surrounding each time step *t*_0_ (blue shaded area), from which the Fourier transform was computed. A 1*/ f* function was fitted to the power spectrum, and removed to obtain the residual power spectrum around *t*_0_. Then, the data were aggregated into 2-Hz bands and normalized, to obtain a plot like in the upper-right, in which the value at each element is a residual power value for in 2-Hz frequency bands at each time step. **Panel (c)**: The covariance matrix of the frequency bands was eigendecomposed using principal component analysis (PCA). Utilizing the first two components per brain region led to a six dimensional *phase-space* space, which we subdivided into small grid cells. The multichannel spectra at each time point was localized to a cell in this space, and we computed the conditional probability of traveling from one cell to another. The Louvain method applied to the network spanned by these conditional probabilities produces clusters, defined as groups of cells where the system is likely to remain within for some time.

The mice’ real-time position was continuously monitored via a webcam sampled at 24 Hz and synchronized with the electrophysiological data. Video data were processed in DeepLabCut [21], a software package for markerless pose estimation based on convolutional deep neural networks. 200 randomly selected frames were hand-labeled for the left ear, right ear, nose, and the beginning of the tail. The corners of the objects were also labeled. We used the ResNet-101 network with 200,000 iterations, and visual inspection was used to confirm accuracy of the marker labels in test frames.

Each video frame was given one of three labels with a corresponding variable value *ζ*: *non-exploration* when all of the mouse’s body markers were outside the boundaries of the object (*ζ*= 1); *exploration* when any of the mouse’s head markers were inside the polygon derived from the object corners of the objects (*ζ*= 2); *novel exploration* when any of the head markers were inside the polygon of the novel object during the test phase (orange dot in Figure 1a) (*ζ*= 3). Exploring the familiar object in the test session was labeled *ζ*= 2.

### 2.2. Data processing and analysis

Electrophysiology data were acquired using Open Ephys hardware with a sampling rate of 30 kHz. Offline, data were imported into MATLAB, down-sampled to 1000 Hz, high-pass filtered at 0.5 Hz, and locally referenced to the average signal from each region. This re-referencing ensured that the signals were locally generated and not volume-conducted from distal brain regions or from the online reference electrode in the skull on top of the cerebellum. The EEGlab toolbox [22] was used for visual inspection of data quality, and for removing non-neural artifacts via independent component analysis using the jade algorithm [23], as we have described previously [19]

### 2.3. The PLC method

The overall aim of PLC is to characterize how multivariate brain activity clusters over time. This involves three steps that are detailed below: (1) Preparing the data via spectral time series decomposition, (2) reducing the dimensionality via PCA, and (3) identifying clusters from time-varying trajectories in the low-dimensional subspace via the Louvain method.

All analyses were implemented in Python 3.7.6 using custom-written code that relied on standard Python libraries like numpy, scipy, pandas and matplotlib, and in particular the python-louvain package, which used for the clustering (https://github.com/taynaud/python-louvain, based on [14]). Code will be made available upon acceptance. Below, we continue with a more detailed step by step discussion of the methods.

For the second step, we reduced the dimensionality of these 50 series via PCA, retaining two components per brain region. In total, this produced six principal component time series. The third step involved clustering the 6D PCA space via Louvain method. The clusters represent specific combinations of the activation of frequency bands in the three brain regions that remain pseudo-invariant for some time, which may point to the brain to be in a certain state. In light to the experiment, we will relate these clusters to exploration behavior of the mice.

#### 2.3.1. Spectral time series decomposition

Data preparation involved obtaining a standardized time series of the relative powers of 2 Hz frequency bands from 0 to 100 Hz in 50 steps (Fig. 1b). We began by segmenting the data into 1-second epochs and applied the following procedure:

- Compute the Discrete Fourier Transform (DFT) by applying Fast Fourier Transform (FFT) to the data in the 1-second segment.
- Fit the function *ρ*(*f*) = *a f* ^*b*^ to the power spectrum, and compute the residual power spectrum after removing this best-fit line. This removes the ‘1*/ f* ‘ component of the spectrum, and thus allows our method to leverage the entire spectrum, rather than being biased by increased energy at low frequencies.
- Crop the resulting ‘residual’ power spectrum of this window between the frequencies of 0 and 100 Hz (excluding 0 itself) and average the values into bands of 2 Hz (resulting in 50 values over the entire frequency range).

The above procedure produced a matrix of time-by-frequency-by-electrodes (per mouse). Next, we averaged the time-by-frequency values across all channels within each brain region, and obtained an estimate of the dominant (“1*/ f* -residualized”) spectral dynamics within each brain region. To remove any biases of certain frequency bands having systematically higher power or power-variance, we z-scored the time series for each frequency.

#### 2.3.2. Principal Component Analysis

We next applied PCA on the residual power spectra over time to obtain a reduced-dimensional representation of the spectral dynamics. We performed the PCA on the time-by-frequency matrices, separately per brain region. The resulting PCs (spectral modes) reflect linear combinations of power across frequencies that maximize the variance of their weighted combinations.

In practice, PCA is obtained as the eigendecomposition of the data covariance matrix (Fig. 1c). In order to get an aggregated result across the 11 mice, we computed the 50-by-50 average covariance matrix of the time series derived above, per brain region. Since the average covariance matrix is symmetric, the PCs constitute an orthonormal basis. The number of PCs to preserve for analyses was selected based on visual inspection of the eigenvalue spectrum (scree plot), which encodes the amount of variance explained by each PC.

Note that although we performed the PCA separately for each brain region, the data for each brain region were acquired simultaneously. This allowed us to localize the data to a reduced PC space that spans all three regions. Using these different dimensions, we constructed a 6D space with the dimensions being (*PC*1_HIP_, *PC*2_HIP_, *PC*1_PAR_, *PC*2_PAR_, *PC*1_PFC_, *PC*2_PFC_). We refer to this space as the *phase-space*. When the data for each mouse at every time point are projected onto this six-dimensional phase-space, it creates a brain state point corresponding to that time point in the phase space. Connecting these points sequentially in time for a specific mouse then produces the trajectory for the state of its brain in the phase-space.

#### 2.3.3. Louvain clustering

Louvain clustering [14] identifies quasi-stationary pockets of the phase-space, in other words, *clusters* in which the data trajectory remains roughly stable for some period of time. Louvain clustering is a graph clustering method and is based on the concept of modularity, which measures the density of links within the clusters with respect to the density of links between clusters. Weights of links are taken into account in the definition of modularity. The method can be applied to find clusters based on the trajectories in the 6D phase-space by discretizing the space into cells. In particular, we gridded the 6D space into one million cells, with 9 cells in each dimension, equally spread in the domain [− 12, 12] for all brain regions – HIP, PFC and PAR. (The value 12 as domain boundaries corresponds to the maximum absolute value of the PCs for the entire time series across all brain regions, which is just above 11.5.) We then constructed a matrix *M* with the following entries

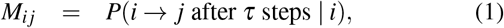

which denotes the conditional probability of the trajectory to move from cell *i* to cell *j* after time *τ*. Here, we used *τ* = 30 ms. The matrix *M* is the transfer matrix, and can be used to statistically simulate the time evolution of the state of the mouse brain throughout the phase space [13]. It can also be seen as a directed network with weighted edges denoting the conditional probability of transfer from one node to another in a given lag time: nodes in the network are then the grid cells in the phase space. Stated differently, Louvain clustering provides an algorithm to identify clusters of grid cells in the 6D phase space in which the mouse brain is (statistically) likely to remain, within for times of order *τ*. Prior to Louvain clustering, we smoothed the time series data using a Gaussian with a full-width-half-maximum of 30-ms (selected to match *τ*).

*τ* is a parameter of the analysis. In initial explorations with pilot data, we found that modularity generally and smoothly decreased with increasing *τ*. Furthermore, the locations and boundaries of clusters were qualitatively comparable with *τ*’s ranging from 5-300 ms. Here we selected *τ* = 30 based on the approximate time windows of spike smoothing [24]. On the other hand, we acknowledge that this parameter selection is somewhat arbitrary, and it is possible that different features of the data would be highlighted by different values of this parameter.

#### 2.3.4. Cluster features

In order to characterize the Louvain clusters and relate them to mouse behavior, we computed the following attributes for each cluster. Some of these features can also be calculated per mouse, but we computed them as aggregated over all datasets.

- *Exploration bias*, a normalized measure of the percentage of time the mice spent exploring the novel object while the brain was in each cluster. Comparing the exploration percentage while in cluster *i* to the overall exploration percentage yielded its *exploration bias*, defined as

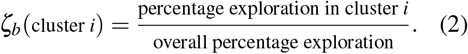

If *ζ*_*b*_(cluster *i*) *>* 1, then the mice explored the object for longer while in brain-state cluster *i* than they do on average over the whole phase-space. If this number is significantly higher than 1 across the group of mice, the mice explore more often than on average when their brain activity’s position in the phase-space places them in cluster *i*, which, via the eigenvectors shown in Fig. 2, can be reconstructed as certain combinations of residual spectra being activated or covarying during exploration.
- *Absolute exploration bias*, defined as *ζ*_abs_ = abs(*ζ*_*b*_− 1). This is the distance of *ζ*_*b*_ from one, and depicts the general strength of the bias (exploration or non-exploration).
- *Average residence time* of state of the mouse’s brain within each cluster. Note that since we are working in 6D space, a cell in a given cluster might share boundaries with cells that do not belong to the same cluster. This results in dramatically reduced residence times due to spuriously frequent entries into and exits from the clusters (stated differently, longer residence periods in the clusters are generally interrupted by this spuriousness). We therefore introduced an additional 30-ms [aligned with the *τ* in Eq. (1)] allowance period to keep count of the residence time if the state of the brain exits and enters a cluster within this time frame. We also omit residence times of 1 or 2 ms in this statistic.
- *Interregionality*, which indicates whether the cluster has more variance in some brain regions compared to others. This indicates whether the cluster is dominated by one region, or whether it comprises dynamics from two or three regions. We defined interregionality (IR) of cluster *i* as 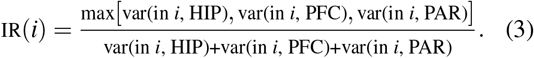

Note that IR ∈[1*/*3, 1]; the lower limit 1*/*3 corresponds to the case where the variance is equally spread across all three brain regions, whereas IR=1 indicates that the variance is exclusively concentrated within one brain region.
- *Mouse specificity*, which indicates whether a cluster is defined from one or a small number of mice, vs. whether data from most or all mice contributed an equal number of data points. Mouse specificity was computed by first calculating the percentage *p*_*m*_(*i*) of cluster *i*’s data points that are from each of the 11 mice *m*. Then the mouse specificity is defined as 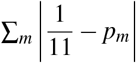 i.e., as the sum of the absolute differences from 1*/*11.
- *Cluster magnitude*, the average distance of data points in the cluster to the origin of the 6D PC space. Distance to the origin was measured by computing the 6D Euclidean distance, weighing each dimension equally.

### 2.4. Statistics

Statistical robustness was checked in multiple respects. To check whether random exploration data can reproduce the same *ζ*_*b*_ distribution across the clusters, the behavioral time series of exploration were shuffled (without changing the brain or PC data), and the above metrics were recomputed. The shape and number of clusters themselves was checked against random permutations by shuffling the PC data, while preserving the internal 6D coordinates (i.e., only shuffling the time sequence of these 6D data points). A final statistical check was the universality of the results across the mice. To this end, we determined *ζ*_*b*_ for each of the eleven mice and performed a Wilcoxon signed rank test. The Wilcoxon test determines whether two samples 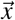 and 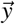 come from the same distribution by checking that the differences 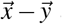 are symmetric around zero. It is different from the regular student’s *t*-test in the sense that the Wilcoxon test is non-parametric — it focuses on the signed ranks, rather than the actual values of the differences. In our case, we used it to test the hypothesis that the novel exploration was consistently in the same clusters across animals.

## 3. Results

### 3.1. General

Across all eleven mice, the average percentage of time spent exploring the novel object was 16.4%. The individual numbers were 15%, 9%, 16%, 9%, 23%, 16%, 13%, 16%, 13%, 37% and 14%.

The parameters *a* (the scaling factor) and *b* (the exponent) of the fits subtracted from the power spectra in each time step were *a* = 0.01± 0.004 and *b* = − 0.36± 0.09. The averages and standard deviations were calculated over all channels, mice, brain regions and parts of the experiment (training + test).

### 3.2. PCA eigenvectors

Inspection of the eigenvalue spectra from the PCA on the residual time series per brain region and per mouse indicates that a large amount of variance in each dataset can be explained only by two PCs (Figure 2). In the interest of convenience and comparability, upon visual inspection, we retained only the top two PCs from each brain region per mouse.

**Figure 2.**
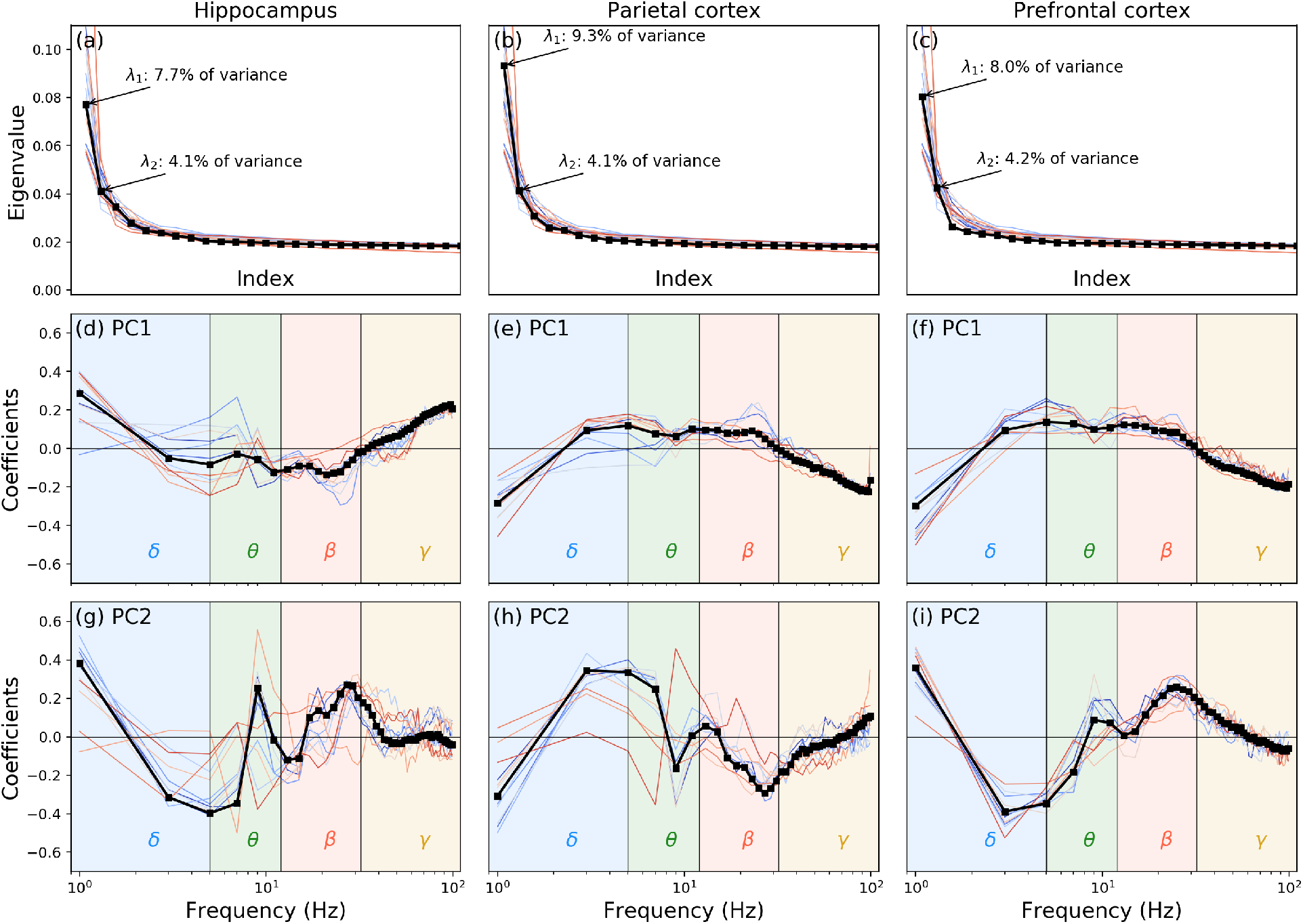
**Panels (a)-(c)**: eigenspectra of the three brain regions. **Panels (d)-(i)**: the top two PCA eigenvectors. For all panels: black indicates those obtained from the average covariance matrix across all eleven mice, while coloured lines indicate individual mice. The background shading in the bottom two rows indicates approximate canonical spectral boundaries for *δ*-, *θ* -, *β* - and *γ*-bands. These bands are purely for reference; the analyses was based on the full spectrum up to 100 Hz, not on individual bands.

Remarkably, the PC eigenvectors for each mouse exhibited strong consistency across the mice, as seen in Fig. 2. This motivated us to compute the average covariance matrix for all the mice together, and then rerun the PCA on this average matrix. The resulting PC eigenvectors are displayed in the same figure in black. We used the latter as the reference set of axes onto which we projected the brain state data for all mice. The advantage of this approach is that it provides us with a “universal” phase space for performing our analyses.

### 3.3. Clusters

Projecting the brain state data onto the 6D (*PC*1_HIP_, *PC*2_HIP_, *PC*1_PAR_, *PC*2_PAR_, *PC*1_PFC_, *PC*2_PFC_) phase-space produces a time series of the 6D coordinates per mouse. In total, data from eleven mice produced 13,644,532 data points. Gridding the 6D space produced 2417 non-empty cells (c.f. the maximum possible 1,000,000 cells). In the calculation of the conditional probabilities for the brain state to transfer from one cell to another, used as entries in matrix *M*, we made sure that only movements between cells within the data of a single mouse are used. The Louvain clustering on matrix *M* identified 60 clusters.

An example cluster is visualized in Fig. 3. The axes are from the first PC of each brain region. The blue colors illustrate the distribution of all data on these axes, and the red colors illustrate the density of the example cluster. Note that the data have higher concentration towards the origin of the phase space. This particular cluster is index #4, contains 273 cells, and comprises 529,167 data points (about 4% of the data).

**Figure 3.**
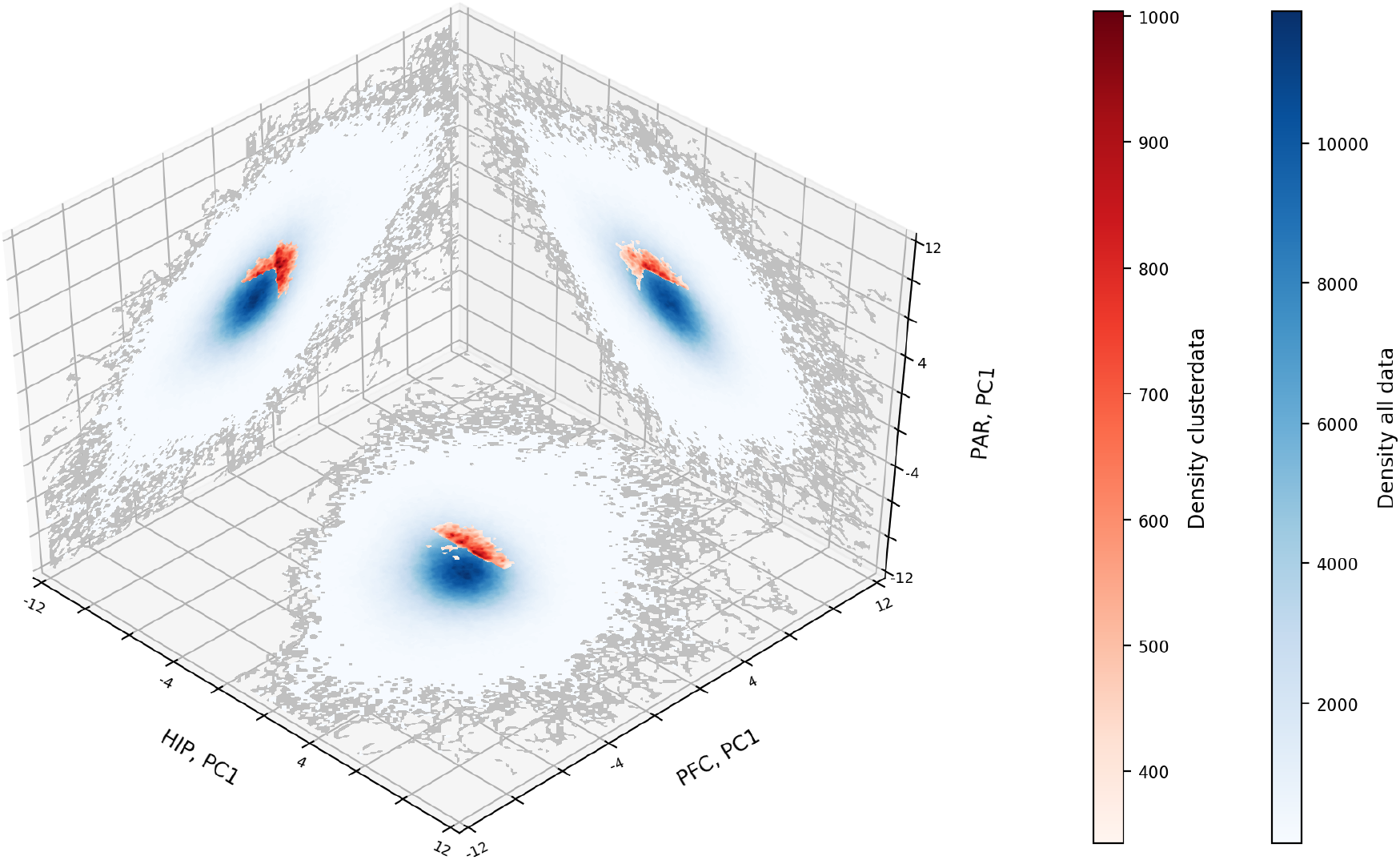
Example cluster found by applying Louvain clustering to the matrix *M*. The axes are the first PCs from each region. In other words, this shows only 3 of the 6 dimensions of the PC space. Blue indicates the distribution of all data, red indicates the distribution of the data of one example cluster. For visualization purposes, the outer edges of the data clouds are marked in gray. In the background, the resolution of the Louvain clustering is marked (9×9).

### 3.4. Cluster attributes

We start investigating the general properties of these clusters by computing the metrics discussed in section 2.3.4. Fig. 4a shows the correlation matrix (Spearman) for the cluster metrics described in the Methods section. We focus on the exploration bias and the absolute bias, because these two metrics are linked to the animals’ behavior. Exploration bias was largely independent of the other metrics.

**Figure 4.**
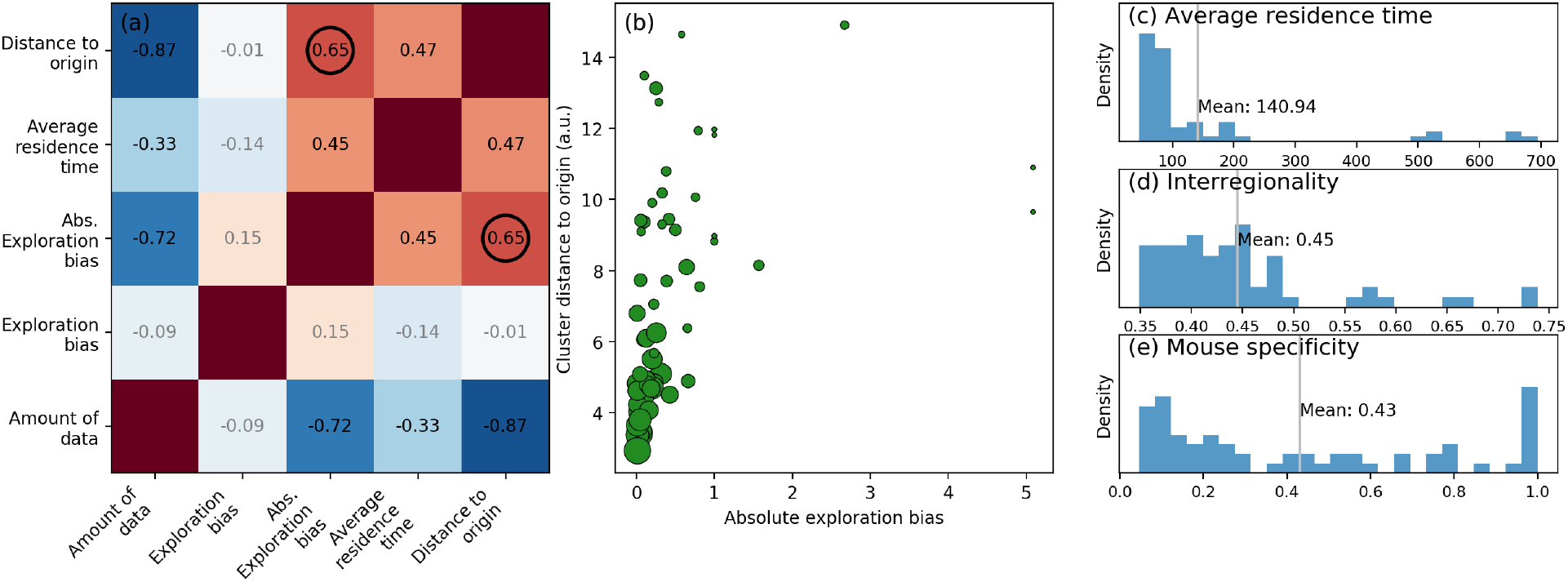
**Panel (a)**: Spearman’s correlation matrix of relevant variables. Negative correlations are denoted in blue, positive correlations are denoted in red. Numerical values of correlations are shown in black for those with correlation significance level *p <* 0.05, other (non-significant) correlations are shown in gray. **Panel (b)**: Cluster distance to origin versus absolute exploration bias, corresponding to the encircled Spearman correlation of 0.6 in panel (a). Scatter marker size in panel (b) corresponds to amount of data points. For reference, we denoted two clusters with their respective amount of data points. **Panel (c)-(e)**: Histograms and associated mean values of (c) average residence time (in ms), (d) interregionality (between 0.33 and 1, higher means more variance in a particular brain region) and (e) mouse specificity (between 0 and 1, higher means more dominated by a smaller set of mice).

Absolute exploration bias, however, was negatively correlated with the amount of data: clusters with more data had exploration biases closer to 1 (see also Fig. 4b).

This is interesting to relate to the positive correlations between absolute exploration bias and residence time and distance to the origin: Clusters in which trajectories remained for longer periods of time — and clusters that were further away from the origin of the PC phase-space — were more likely to be modulated by behavior. One can think of these “peripheral” clusters are requiring more energy in the PC space to push away from the origin, and may therefore reflect brain states that deviate from the “status-quo”.

In contrast, the clusters located towards the center of the PC space contain much more data per cluster, and also more transitions across clusters.

Fig. 4c-e shows distributions for three of the cluster properties. The residence times ranged from tens of ms to 700 ms, but were concentrated around 100 ms, with an average of 140 ms. This is consistent with findings from other studies showing that neural “microstates” tend to last 50-150 ms [25, 5, 26]. Of particular importance is the clustering of interregionality and mouse specificity towards smaller numbers – these two panels illustrate that most clusters comprised data from all three brain regions and from most animals. Thus, the clusters are defined by dynamics that are shared across the animals and reflect coordination across the brain.

### 3.5. Exploration data and Wilcoxon test

To investigate the significance of the exploration bias, we computed *ζ*_*b*_ for all clusters, both for individual mice and for the group-aggregated data. Results are shown in Fig. 5a (squares for individual data and circles for aggregated data). We sorted the clusters by *ζ*_*b*_ of the full data set. Around half of the clusters had below-average exploration bias. Most clusters showed individual variability across animals (note that the aggregated results can be considered a weighted average, because not all animals had the same amount of data per cluster). Panel (c) shows that clusters also varied in the total amount of data per cluster, with the clusters exhibiting strong exploration biases having overall less data.

**Figure 5.**
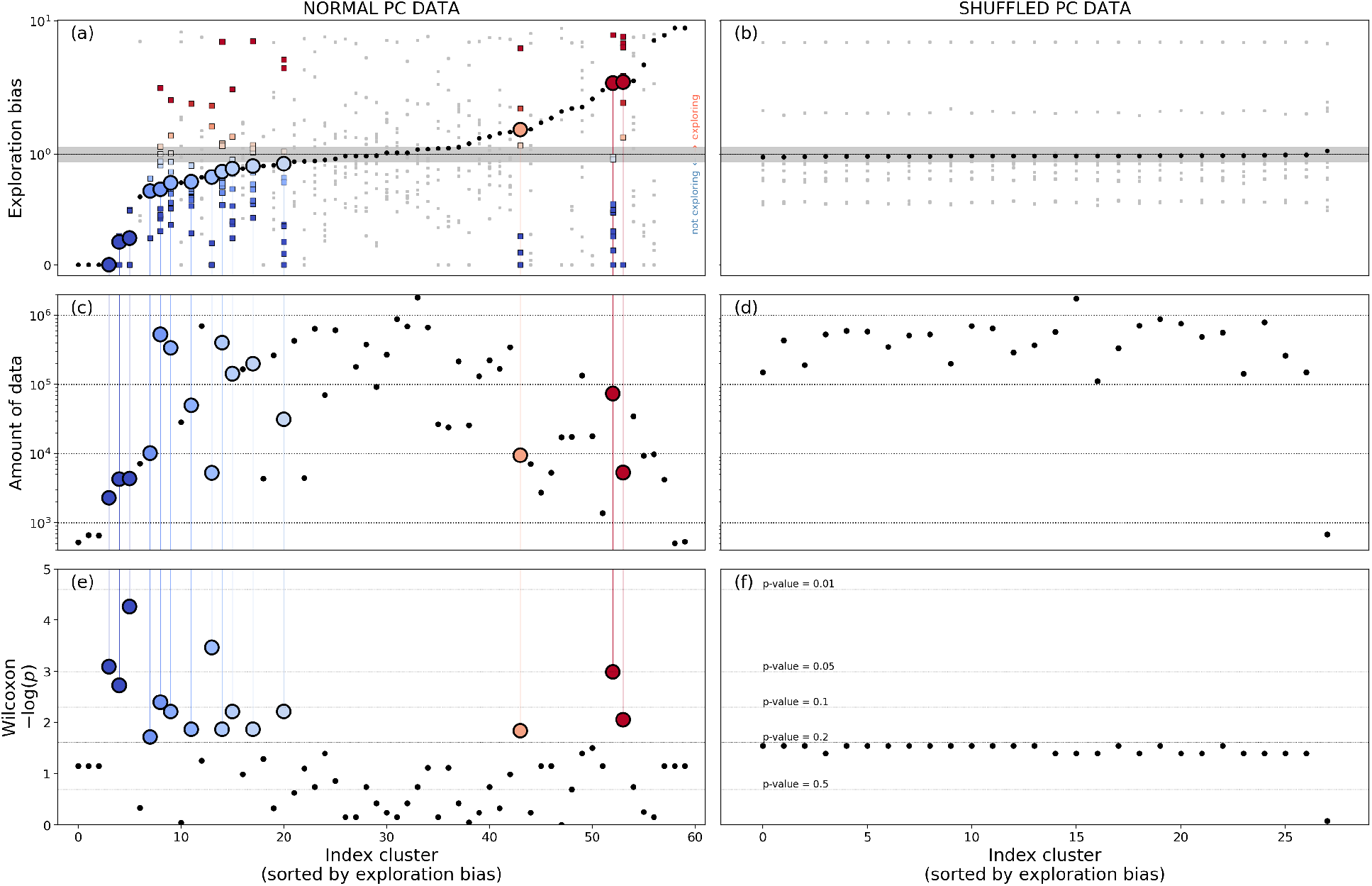
Attributes of the PC clustering (left column) and the clustering applied to shuffled PC data (right column). Exploration bias for every cluster (panels a-b) is compared to the cluster’s respective data amount (panels c-d) and the Wilcoxon *p*-values, that test the consistency of exploration bias values across the 11 animals (panels e-f). Clusters with *p <* 0.2 on the left hand side are highlighted, all other clusters are denoted by black dots. Squares in panels a-b (Colored and gray) show the respective individual biases per mouse. Grey shaded areas in panels a-b indicate the estimated exploration bias range (mean +/- 2*σ*) among 100 times shuffling the exploration data without changing the LFP data.

We performed three sets of statistical evaluations on the exploration bias results. First, we tested the individual data against a bias score of 1 (corresponding to no exploration bias), which tests the consistency of the sign of the bias across animals. Only animals with at least 2% of their data in the particular cluster are used in this analysis, as accounting for mice with smaller amounts of data may results in non-representative values of the exploration bias in these clusters. Results are shown in panel (e), with circles indicating data points of *p <* 0.2. Four clusters exceeded a *p <* .05 threshold. This result is consistent with the cross-animal variability in bias scores seen in panel (a).

The second test was based on shuffling the data to determine whether our results could have been observed in random data in the same reduced PC space. To this end, we scrambled the PC time series (which shuffles the links between grid cells), a re-applied Louvain clustering. Results are shown in Fig. 5a,d,f. Here we observed only 28 clusters with uniform amount of data and clustering bias.

The final set of statistics involved shuffling the temporal mapping between behavior and brain. We cut the behavior time series at a random time point and swapped the second for the first segment, and then recomputed *ζ*_*b*_ for each cluster. Note that this procedure preserves both the LFP data and the behavior data, only randomizing their temporal mapping. This procedure was repeated 100 times, and the gray shaded region in Fig. 5a illustrates two standard deviations around the mean shuffled exploration bias.

Overall, these tests confirm that the clustering structure found in the data was not observed when clustering noise, and that there is individual variability in the specific clusters in which animals investigated the novel object that is not accounted for by random reshuffling of the behavioral data.

## 4. Discussion

### 4.1. Multivariate neural dynamics

Neural activity is diverse and dynamic over space, time, and frequency [27, 9, 2]. Characterizing these dynamics remains a major challenge in neuroscience. Here we adopted a recently developed method [13, 17] to characterize phase-space trajectories in multi-site LFP recordings in mice. An advantage of PLC is that it allows for a mix of linear (PCA) and nonlinear (Louvain clustering) methods to identify trajectory-based clustering in a reduced-dimensional space.

### 4.2. Consistency of PCs across mice

It is striking that the eigenvectors of the top two PCs were so similar across the 11 different mice. This is not a trivial result due to biased selection, because we simply took the top two components rather than, e.g., picking the components that maximized a cross-mouse correlation. It is also not a trivial result of the 1*/ f* -like nature of the power spectrum, because this feature was removed prior to the covariance matrix generation.

Instead, we interpret this result to indicate that there are fundamental characteristics of this cortical-hippocampal network that are (1) concentrated in a lower-dimensional space, and (2) conserved across mice. On the other hand, the PCs were not fully identical across mice, and the residual variance likely comprises a combination of unique factors and sampling variability.

Whether to apply multivariate decompositions to data pooled over individuals is debated in the neuroscience literature [28, 29, 30], and leads to a trade-off between increased generalization vs. increased sensitivity to individuals.

We decided to use the same PC space for all mice in order to have a “universal” framework, which facilitated pooling the cluster characteristics across all mice. The mouse specificity in Fig. 4e indicates that there are many clusters in which most or all mice are well represented — indicating that inter-regional brain dynamics of distinct mice are situated in the same clusters — and there were also several clusters with near-1 values of mouse specificity, indicating that some clusters were unique to individual mice.

### 4.3. Interpretation of clusters and usage of Louvain clustering

Over time, the trajectory of the brain state moves around in the 6D PC space. These trajectories are not random, and instead remain within relatively confined “provinces” for periods of time. Louvain clustering identifies these clusters. A key parameter of Louvain clustering is *τ*, which is the time-scale at which a trajectory is considered stationary. In our pilot analyses with data from one mouse (not shown here), we found that the clustering was stable for a range of *τ* values, and thus we selected *τ* = 30 ms in the interest of consistency. It is possible that different features of the data would be highlighted for larger differences in this parameter, e.g., over hundreds of ms or seconds. Determining how trajectories and clustering are related across temporal scales is an interesting avenue for future research to explore.

Given that the PLC method is several steps away from the raw data, one might wonder whether these clusters are meaningful, or simply reflect residual noise. We addressed this in two ways. First, Louvain clustering applied to shuffled data showed qualitatively distinct patterns of results compared to the real data (e.g., Figure 4). Indeed, clustering shuffled data produced only 28 clusters, compared to the 60 clusters in the real data, even though the amount of data was the same.

Second, we examined the relationship between the cluster characteristics and exploration behavior. The results shown in Figs. 5 and 4 suggest that clusters close to the center of the PC phase-space were generally not modulated by behavior, whereas clusters further away from the origin had exploration biases that deviated from 1. These findings suggest that the brain space trajectories associated with active behavior are “high-energy” states that are maintained for relatively longer periods of time.

On the other hand, there was also considerable individual variability in the exact exploration bias scores over the different animals, evidenced by the relatively large p-values in the Wilcoxon test (although one must keep in mind that with N=11, the Wilcoxon test can only return a p<.05 result if 10/11 datapoints have the same sign). Some clusters had relatively little or no data for some animals. This finding raises the question of whether the clustering should have been done on individual animals instead of on the group data. As we wrote earlier, group-defined clusters have several advantages in terms of comparability of cluster findings and “universality” of the spectral-temporal structure of the data. Nonetheless, these are analysis choices that users can determine on their own; our primary purpose here was to illustrate the utility and interpretability of the PLC method.

The quadratic relationship between exploration bias and distance to the phase-space origin is highlighted by the significant correlation with *absolute* exploration bias. Thus, although the direction of the exploration bias score varies across animals and clusters, there is in general more behavioral variability as the data move towards the periphery of the phase-space. It is possible that the center of the space reflects brain states of relatively low energy or of transitions between states of higher energy. On the other hand, one must keep in mind that the outer edges of the phase-space are relatively sparse, which means that the clusters had overall less data. This can become statistically problematic in the extreme case of, for example, a cluster comprising only 50 data points from a single mouse that is exploring the object for the entire time window.

### 4.4. Limitations and future directions

The PLC method is not without limitations. As mentioned above, clusters with relatively little data, or that are dominated by a single animal, can provide cluster characteristics that may not be fully representative of group dynamics. In the present application, the higher density of data towards the center of the 6D phase-space means that clusters at the outer edges of the data cloud have overall less data. This is apparent in Fig. 5c, which shows differences in densities of clusters spanning four orders of magnitudes. We chose not to remove any clusters based on total amount of data, although we did exclude animals from group-level t-tests if those animals had too little data in that cluster.

A second limitation is the operational definition of a “cluster” in noisy data such as LFP. Small and fast jumps out of, and then back into, a cluster may reflect noise or a true rapid fluctuation between states. This would manifest as low residence times. We chose to filter out jump-but-return trajectories that were faster than 30 ms.

Third, the physiological meaning of PLC-derived clusters remains unknown. We speculate that the brain remains in a particular “state” while in these clusters, similar to the interpretation of EEG microstates [25, 5, 26]. This could be investigated more in the future, for example by investigating spiking dynamics in different LFP-defined clusters.

The PLC method is general, and can be applied to many additional tasks or behavior elements. It can also be applied to larger-scale measurements such as EEG and MEG. Depending on the application and the data, it may be the case that the phase-space dimensions have to be adapted, to, for example, focus on a single brain region, or more or fewer principal components.

## Acknowledgements

We thank Mihaela Gerova for assistance with data preparation and video processing with DeepLabCut. MXC, ASCF, and this research were funded by an ERC-StG 638589 and a Hypatia fellowship from the Radboud University Medical Center.

## Author contributions statement

MD, DP and MXC conceived the method. AF performed the experiments. MD developed the code and ran the analysis. MD wrote the first draft of the manuscript. All authors contributed to the final manuscript.

## Additional information

### Accession codes

Code and data will be available upon publication at data.donders.ru.nl.

